# Advances in Gene Ontology Utilization Improve Statistical Power of Annotation Enrichment

**DOI:** 10.1101/419085

**Authors:** Eugene W. Hinderer, Robert M. Flight, Rashmi Dubey, James N. MacLeod, Hunter N.B. Moseley

## Abstract

Gene-annotation enrichment is a common method for utilizing ontology-based annotations in these gene and gene-product centric knowledgebases. Effective utilization of these annotations requires inferring semantic linkages by tracing paths through the ontology through edges in the ontological graph, referred to as relations. However, some relations are semantically problematic with respect to scope, necessitating their omission lest erroneous term mappings occur. To address these issues, we present GOcats, a novel tool that organizes the Gene Ontology (GO) into subgraphs representing user-defined concepts, while ensuring that all appropriate relations are congruent with respect to scoping semantics. Here, we demonstrate the improvements in annotation enrichment by re-interpreting edges that would otherwise be omitted by traditional ancestor path-tracing methods.

We demonstrate that GOcats’ unique handling of relations improves enrichment over conventional methods in the analysis of two different gene-expression datasets: a breast cancer microarray dataset and several horse cartilage development RNAseq datasets. With the breast cancer microarray dataset, we observed significant improvement (one-sided binomial test p-value=1.86E-25) in 182 of 217 significantly enriched GO terms identified from the conventional path traversal method when GOcats’ path traversal was used. We also found new significantly enriched terms using GOcats, whose biological relevancy has been experimentally demonstrated elsewhere. Likewise, on the horse RNAseq datasets, we observed a significant improvement in GO term enrichment when using GOcat’s path traversal: one-sided binomial test p-values range from 1.32E-03 to 2.58E-44.

## Introduction

### Ontologies and gene set enrichment analyses

Biological and biomedical ontologies such as Gene Ontology (GO) [1] are indispensable tools for systematically annotating genes and gene products using a consistent set of annotation terms. Ontologies are used to document new knowledge gleaned from nearly every facet of biological and biomedical research today, from classic biochemical experiments elucidating specific molecular players in disease processes to omics-level experiments providing systemic information on tissue-specific gene regulation. These ontologies are created, maintained, and extended by experts with the goal of providing a unified annotation scheme that is readable by humans and machines [2]. With the advent of transcriptomics technologies, high-throughput investigation of the functional impact of gene expression in biological and disease processes in the form of gene set enrichment analyses represents one important use of GO [3]. Many different tools exist to utilize GO annotations in enrichment analyses [4–6]. However, all current methods fail to utilize all the semantic information available in this ontology due to inconvenient features in the anatomy of GO.

### Anatomy of the Gene Ontology

The GO database represents a controlled vocabulary (CV) of biological and biochemical terms that are each assigned a unique alphanumeric code, which is used to annotate genes and gene products in many other databases, including UniProt [7] and Ensembl [8]. The ontology is divided into three sub-ontologies: Cellular Component (CC), Molecular Function (MF), and Biological Process (BP). Each can be envisioned as a graph or network where terms are nodes connected by edges, referred to as relations, that describe how each term relates to one another. For example, the term “DNA methylation” (GO:0006306) is connected to the term “macromolecule methylation” (GO:0043414) by the is_a relation. In this case, ontological terminology defines the term “macromolecule methylation” as a “parent” of the term “DNA methylation.” The three sub-ontologies mentioned are “is_a disjoint” meaning that there are no is_a relations connecting any node among the three ontologies. However, other relations, such as “regulates,” connect nodes of separate sub-ontologies. Relations of interest to this study are part_of and has_part. These are like is_a in that they describe scope, i.e. relative generality or encompassment, but are separate in that is_a represents true sub-classing of terminology while part_of and has_part describe part-whole (mereological) correspondence. Therefore, we consider scoping relations to be comprised of is_a, part_of, and has_part, and mereological relations to be comprised of part_of and has_part.

There are three versions of the GO database, each containing aspects of the CV with varying complexity: *go-basic* is filtered to exclude relations that span across multiple sub-ontologies and to include only relations that point toward the root of the ontology; *go* or *go-core* contains additional relations, such as has_part that may span sub-ontologies and which point both toward and away from the root of the ontology; and *go-plus* contains yet more relations in addition to cross-references to entries in external databases like the Chemical Entities of Biological Interest (ChEBI) ontology [9]. The first and second versions are available in the Open Biomedical Ontology (OBO) flat text file formatting, while the third is available only in the Web Ontology Language (OWL) RDF/XML format.

### Path traversal issues in GO

Ontological graphs are typically designed as directed graphs, meaning that every edge has directionality, or directed acyclic graphs (DAGs), meaning that no path exists that leads back to a node already visited if one were to traverse the graph stepwise. This allows the graph to form a complex semantic model of biology containing both general concepts and more-specific (fine-grained) concepts. The “parent-child” relation hierarchy allows biological entities to be annotated at any level of specificity (granularity) with a single term code, as fine-grained terms intrinsically capture the meaning of every one of its parent and ancestor terms through the linking of relation-defining is_a edges in the graph. However, it is deceptively non-trivial to reverse the logic and organize similar fine-grained terms into general categories—such as those describing whole organelles or concepts like “DNA repair” and “kinase activity”—without significant manual intervention. This is due, in part, to the lack of explicit scoping, scaling, and other semantic correspondence classifiers in relations; it is not readily clear how to classify terms connected by non-is_a relation edges. Although edges are directional, the semantic correspondence between terms connected by a scoping relation is computationally ambiguous, e.g. assessing whether term 1 is more/less general or equal in semantic scope with respect to term 2 is currently not possible without explicitly defining rules for such situations.

Ambiguity in assessing which term is more general in a pair of terms connected by a relation edge is confounded by the fact that edges describing mereological relations, such as part_of and has_part, are not strictly and universally inverse of one another. For instance, while every “nucleus” is part_of “cell,” not every “cell” has_part “nucleus.” Similarly, while every “nucleus” has_part “chromosome”, not every “chromosome” is part_of “nucleus” under all biological situations. Therefore, mereological edges are not necessarily reciprocal. Ontological logic rules, called axioms, ensure that this logic is maintained in the graph representation by allowing edges of the appropriate type to connect terms only if the inferred relation is universal [10, 11]. This axiomatic representation is crucial to avoid making incorrect logical inferences regarding universality but does nothing to facilitate categorization of terms into parent concepts, especially since some mereological edges point away from the root of the ontology, toward a narrower scope. If these edges are followed, terms of more broad scope may be grouped into terms of more narrow scope, or worse, cycles may emerge which would abolish term hierarchy and make both categorization and semantic inference impossible. To circumvent this problem, some ontologies release versions that do not contain these types of edges. For GO, this is accomplished by go-basic. However, information is lost when these edges are removed from the graph. If attempting to organize fine-grained terms into common concepts using the hierarchical structure, this information loss can be significant because many specific-to-generic term mappings can utilize the same edge in many paths.

### Axiomatic versus semantic scoping interpretation of mereological relations in GO

While ensuring mereological universality in relation associations using current axioms is important within the purview of ontology development, for those interested in organizing datasets of gene annotations into relevant concepts for better interpretation—such is the case in annotation enrichment—it is important to utilize the full extent of the information within an ontology.

Current axiomatic representation of mereological relations requires the use of ontology versions which lack certain relations [12], resulting in a loss of retrievable information. If has_part edges—which point toward terms of narrower scope—were to be inversed to resemble part_of edges—ensuring that all edges point toward terms of a broader scope—terms could be effectively categorized with respect to semantic scope using the native graph hierarchy without losing any information in the process. However, this isn’t logically possible because of issues dealing with universality.

Therefore, we acknowledge the importance of existing axioms which prohibit reversing mereological edges in ontologies under the context of drawing *direct* semantic inferences. However, we maintain that in the context of detecting enriched broad concepts based on “summarizing” annotated fine-grained terms contained within differential annotation datasets, it is appropriate to evaluate mereological relations from a scoping perspective, which requires that all mereological edges point to their whole. This conundrum preventing the comprehensive categorization of GO terms can be dealt with by adding a single new relation to the ontology: part_of_some. Semantically, this relation deals with both the issue of universality and with the issue of the direction of granularity.

### GO Categorization Suite (GOcats)

For the issues stated above, we have developed a new tool called the GO Categorization Suite (GOcats). Fundamental to GOcats’ categorization algorithm is the re-evaluation of the has_part edge as part_of_some—correcting semantic correspondence inferences while ensuring ubiquitous use of all categorization-relevant relations in GO.

In comparing GOcats’ inclusion of re-evaluated has_part relations to the traditional method of ignoring has_part relations altogether and to the erroneous method of misinterpreting native has_part directionality, we illuminate the theoretical extent of information loss or potential for misinterpretation of has_part relations, respectively. Furthermore, in two independent enrichment analyses of real data—from a publicly available breast cancer dataset [13] and from preliminary data investigating horse cartilage development [14], we demonstrate that GOcats’ reinterpretation of has_part can retain all information from GO while drawing appropriate categorical inferences in the context of annotation enrichment. Finally, we show that this reinterpretation has the added benefit of improving the statistical power of annotation enrichment analyses.

## Design and Implementation

The *go-core* version of the GO database was chosen in favor of the *go-basic* version, because it contains the has_part edge relation which points away from the root of the ontology and because it contains other edges which connect the separate subontologies. Since one of our goals is to reinterpret mereological relations with respect to semantic scope, it is necessary that these relations be evaluated. Similarly, we excluded the *go-plus* version from this investigation, because we are not yet concerned with the reevaluation of the additional relations or database cross-references provided by *go-plus*.

While *go-basic* is a true DAG, *go-core* is not strictly acyclic due to the additional has_part relations. However, when we inversed traversal of has_part into the part_of_some interpretation, acyclicity was maintained. Therefore, we refer to our modified *go-core* graph as a DAG. GOcats is a Python package written in version 3.4.2 of the Python program language [15]. GOcats parses go-core and represents it as a DAG hierarchal structure. GOcats extracts subgraphs of the GO DAG (sub-DAGs) and identifies a representative node for each category in question (Figure 1). While GOcats’ categorization algorithms are a major feature of the software, it is not a focus of this study. Full API documentation for GOcats is available online (https://gocats.readthedocs.io).

**Figure 1.**
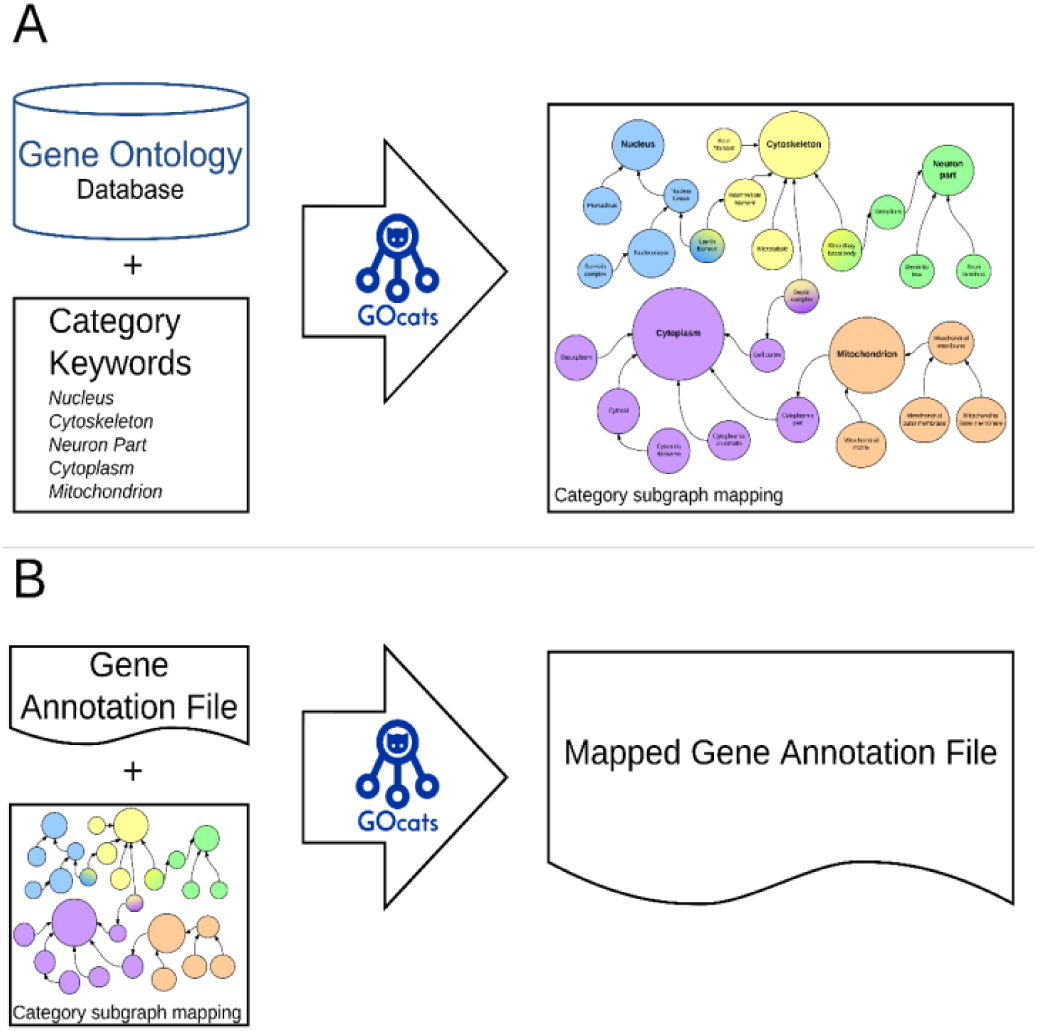
GOcats data flow diagram for creating categories of GO. A) GOcats enables the user to extract subgraphs of GO representing concepts as defined by keywords, each with a root (category-defining) node. B) Subgraphs extracted by GOcats are used to create a mapping from all sub-nodes in a set of subgraphs to their category-defining root node(s). This allows the user to map gene annotations in GAFs to any number of customized categories.

To overcome issues regarding scoping ambiguity among mereological relations, we hard-coded assigned properties indicating which term was broader in scope and which term was narrower in scope to each edge object created from each of the scope-relevant relations in GO. For example, in the node pair connected by a part_of or is_a edge, node 1 is narrower in scope than node 2. Conversely, node 1 is broader in scope than node 2 when connected by a has_part edge (Table 1, Figure 2). This edge is therefore reinterpreted by GOcats as part_of_some. While the default scoping relations in GOcats are is_a, part_of, and has_part, the user has the option to define the scoping relation set. For instance, one can create go-basic-like subgraphs from a go-core version ontology by limiting to only those relations contained in go-basic. For convenience, we have added a command line option, “go-basic-scoping,” which allows only nodes with is_a and part_of relations to be extracted from the graph.

**Table 1.** Prevalence of relations in the Gene Ontology and suggested semantic correspondence classes to reduce ambiguity.

| Relationship | Prevalence in GO (CC+BP+MF) | Prevalence in GO CC | Prevalence in GO BP | Prevalence in GO MF | Correspondence Class | Correspondence Members |
| --- | --- | --- | --- | --- | --- | --- |
| is_a | 72455 | 5591 | 54689 | 12175 | Scoping (hyponymy) | hyponym "is_a" hypernym |
| part_of | 8613 | 1702 | 5751 | 1160 | Scaling (meronymy) | meronym "part_of" holonym |
| has_part | 736 | 156 | 339 | 241 | Scaling (meronymy) | holonym "has_part" meronym |
| happens_during | 24 | 0 | 24 | 0 | Spatiotemporal (process-process) | process "happens_during" process |
| ends_during | 1 | 0 | 1 | 0 | Spatiotemporal (process-process) | process "ends_during" process |
| occurs_in | 181 | 0 | 180 | 1 | Spatiotemporal (process-entity or process-process) | process "occurs_in" entity<br>OR<br>process "occurs_in" process |
| regulates | 3368 | 0 | 3322 | 46 | Active (actor-subject) | actor "regulates" subject |
| positively_regulates | 2916 | 0 | 2880 | 36 | Active (actor-subject) | actor "positively_regulates" subject |
| negatively_regulates | 2937 | 0 | 2285 | 52 | Active (actor-subject) | actor "negatively_regulates" subject |
| regulated_by<sup>‡</sup> | 0 | 0 | 0 | 0 | Active (actor-subject) | subject "regulated_by" actor |
| before<sup>‡</sup> | 0 | 0 | 0 | 0 | Spatiotemporal (prior-latter) | prior "before" latter |

‡ These relationships are not found in GO but are part of the Relations Ontology

**Figure 2.**
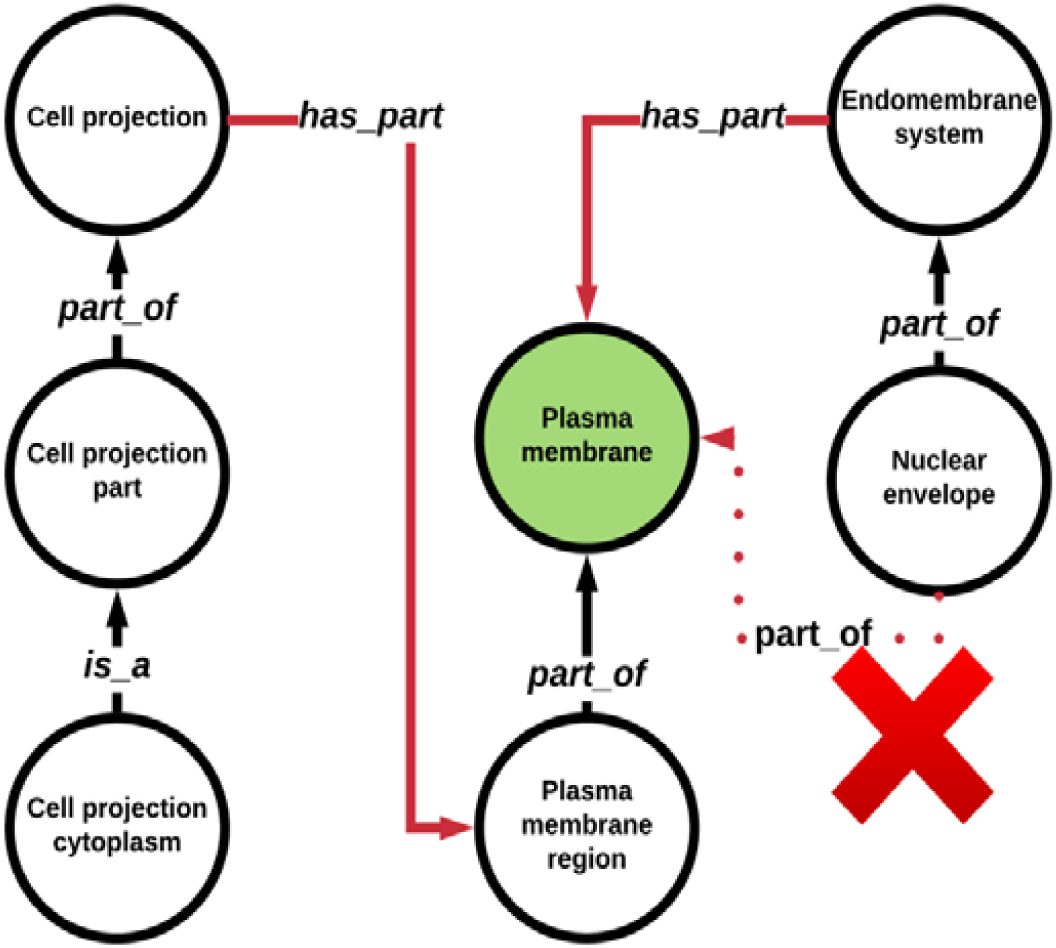
The has_part relation creates incongruent paths with respect to semantic scoping. Some tools may create questionable GO term mappings, i.e. “nuclear envelope” to “plasma membrane,” since the has_part relation edges point in from super-concepts to sub-concepts. GOCats avoids this by re-interpreting the has_part edges into part_of_some edges.

## Results

### GOcats’ reinterpretation of the has_part relation increases the information retrieval from GO and avoids potential misinterpretations of ambiguous relationship inferences

GOcats reevaluates path tracing for the has_part edge to make it congruent with other relations that delineate scope. With path tracing unchanged, has_part edges lead to erroneous term mappings unless they are completely excluded from the ontology. To evaluate the extent of incorrect semantic interpretation conferred by has_part relations, we calculated all potential false mappings (pM_F_) between nodes for a given GO sub-ontology by counting the number of mappings from all children of a has_part edge to all parents of a has_part edge assuming the original GO has_part edge directionality. Next, we compared the pM_F_ to the total number of true mappings (M_T_) for a given GO sub-ontology to evaluate the possible magnitude of their impact (Supplementary Data 1, Equations 1-5, Scripts Repository 1,2). As shown in Table 2, there are 23,640 pM_F_s in Cellular Component, 8,328 pM_F_s in Molecular Function, and 89,815 pM_F_s in Biological Process. Comparatively, the amount of pM_F_s is 42%, 13%, and 16% the size of the M_T_, in Cellular Component, Molecular Function, and Biological Process, respectively.

**Table 2.** Prevalence of potential has_part relation mapping errors in GO.

| Sub-Ontology | Estimated Potential False Mappings (epM<sub>F</sub>) | True Mappings (M<sub>T</sub>) | $M_T \cap epM_F$ | Potential False Mappings $pM_F = epM_F - (M_T \cap epM_F)$ | True Mappings without HP (IA_PO M<sub>T</sub>)* | Lost Mappings (M<sub>T</sub> - IA_PO M<sub>T</sub>)* |
| --- | --- | --- | --- | --- | --- | --- |
| Cellular Component | 30036 | 56025 | 6396 | 23640 | 49679 | 6346 |
| Molecular Function | 10074 | 62436 | 1746 | 8328 | 56194 | 6242 |
| Biological Process | 93092 | 555543 | 3277 | 89815 | 527869 | 27674 |

\* IA\_PO refers to a graph created with only is\_a and part\_of relationship edges.

The conventional solution to avoid these errors is to use versions of ontologies that remove edges like has_part. [16]. Considering the number of possible mappings between terms as a measure of information content, we quantified the loss of information acquired when has_part is omitted during mapping by subtracting the number of M_T_ in graphs containing is_a, part_of, and has_part edges from those with only is_a and part_of edges. As shown in Table 2, Cellular Component lost 6,346 mappings, Molecular Function lost 6,242 mappings, and Biological Process lost 27,674 mappings, which equates to 11%, 10%, and 5% loss of information in these sub-ontologies, respectively. It is important to note here that the mapping combinations were limited to those nodes containing is_a, part_of, and has_part relations only. Because paths in GO are heterogeneous with respect to relation edges, this loss of information is a lower-bound estimate since other relations exist that connect additional nodes erroneously. This is especially true for Biological Process, which has many regulatory relations that were not evaluated here.

While the potential for false mappings are high considering the has_part relation alone, this statistic does not illuminate the scale of the issue facing users of current ontology mapping software. Importantly, it does not address a fundamental limitation and danger facing software like map2slim (M2S) [5], which non-discriminately evaluates relation edges. For example, terms linked by an active relation like *regulates*, or by the has_part edge are categorized as if they are related by a scoping relation like is_a. Therefore, we calculated the total number of possible mappings produced by M2S and enumerated the intersection of these mappings against those made by GOcats which were constrained to paths that contained only scoping relations, is_a, part_of, and has_part (Supplementary Data 2, Equations 6 and 7). Overall, M2S made 325,180 GO term mappings, i.e. categorizations, which did not intersect GOcats’ full set of corrected scoping relation mappings. We consider these false mapping pairs (M_pair, M2S_), since they represent a problematic evaluation of scoping semantics. This contrasted with 710,961 correct mappings that intersected the GOcats mapping pairs (M_pair, GOcats_) giving a percent error of 31.4%. Cellular Component, Molecular Function, and Biological Process contained 22,059, 29,955 and 273,166 erroneous mappings, which accounted for respective percent errors of 30.7%, 34.8%, and 31.1% (Table 3).

**Table 3.** Summary of GO term mapping errors resulting from misevaluation of relations with respect to semantic scoping.

| (Sub) Ontology | Map2Slim Mappings (M<sub>pair,M2S_ont</sub>)* | GOcats Scoping Mappings (M<sub>pair,Gocats_ont</sub>)* | Potentially false Map2Slim Mappings $pM_{F,M2S} = M_{pair,M2S} - (M_{pair,M2S} \cap M_{pair,Gocats\_all})^*$ | Map2Slim Correct Mappings $M_{T,M2S} = M_{pair,M2S} \cap M_{pair,Gocats\_all}^*$ | Possible Map2Slim Error Fraction $pM_{F,M2S} / M_{pair,M2S\_ont}$ |
| --- | --- | --- | --- | --- | --- |
| All GO | 1036141 | 820467 | 325180 | 710961 | 0.314 |
| Cellular Component | 71835 | 56025 | 22059 | 49776 | 0.307 |
| Molecular Function | 86163 | 62436 | 29955 | 56208 | 0.348 |
| Biological Process | 878143 | 555543 | 273166 | 604977 | 0.311 |

\* GOcats\_all refers to GOcats-derived mapping pairs across all of GO, while GOcats\_ont refers to GOcats-derived mapping pairs for the indicated ontology in each row.

### GOcats’ reinterpretation of has_part relations provides improved annotation enrichment statistical power

We incorporated GOcats-derived ontology ancestor paths (paths from fine-grained terms to more general, categorical terms) into the categoryCompare version 1.99.158 [17] annotation enrichment analysis pipeline and performed annotation enrichment on an Affymetrix microarray dataset of ER+ breast cancer cells with and without estrogen exposure [13]. We compared these enrichment results to those produced when unaltered ancestor paths from GO—excluding the has_part relation— were incorporated into the same categoryCompare pipeline (Supplementary Data 3, Scripts Repository 3).

We also performed enrichment analyses comparing the ancestor traversals of DEseq2 differential gene expression datasets across time points during the fetal development of two cartilage tissue types in *Equus caballus* (Supplementary Data 4-5, Scripts Repository 4).

Assessment of adjusted p-values from significantly enriched terms using GOcats’ paths versus the traditional method that omits has_part edges shows that GOcats reliably improves the statistical significance of term enrichment results through its improved re-interpretation of relation semantics (Figure 3, Supplementary Data 6). In the breast cancer dataset, of the 217 significantly enriched terms found using the traditional enrichment method at an alpha of 0.01 for FDR-adjusted p-values, 182 had adjusted p-values that were improved when GOcats part_of_some paths were used. This number of improved p-values is statistically significant as indicated by a one-sided binomial test p-value of 1.86E-25.

**Figure 3.**
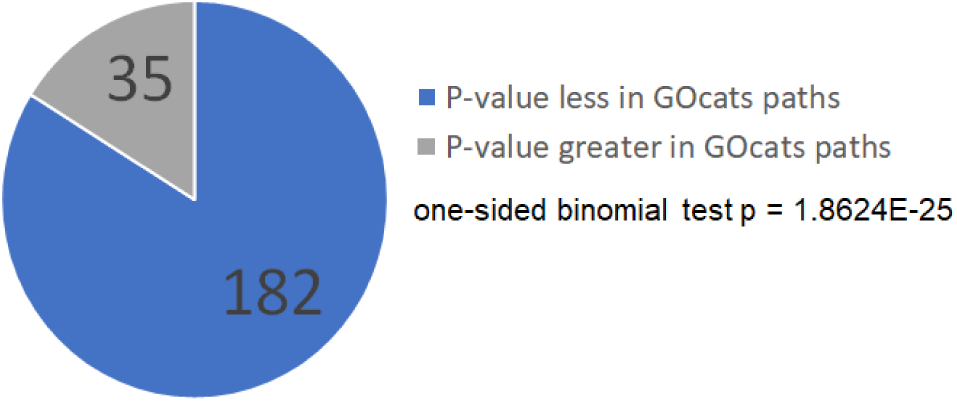
Comparison of adjusted p-values for significantly-enriched annotations using GOcats paths vs excluding has_part edges. Most significantly-enriched GO terms had an improved p-value when GOcats re-evaluated has_part edges for the enrichment of the breast cancer data set in this investigation.

Additionally, GOcats was able to identify 15 unique significantly-enriched terms at an alpha of 0.01 for adjusted p-values that would otherwise be omitted due to the loss of has_part edges (Supplementary Data 7). Four of these terms involve purinergic nucleotide receptor activity, which has been implicated elsewhere in other investigations related to breast cancer [18].

GOcats’ path tracing showed similar improvements when comparing p-values from GO annotation enrichment derived from the differential gene expression analyses between horse cartilage development time points (Table 4). In this analysis (see Supplementary Data 4), neighboring time point analyses (early and late) were compared to extreme time point analyses (extreme). The traditional enrichment method yielded between 82 to 233 total enriched terms, with 67% to 92% of these terms’ adjusted p-values being improved when GOcats ancestor path tracing was used. Quantifying the improvements in the p-values via a binomial test generates p-values ranging from 1.32E-03 to 2.58E-44. All but one of the binomial test p-values was below 6.22E-21; however, the comparison of the fetal interzone tissue fetal at 45 days to neonatal epiphyseal cartilage had drastically fewer total enriched terms. Furthermore, GOcats was able to identify additional significantly-enriched terms from the first and second neighboring time point analyses as compared to the traditional method applied to the extreme analysis. As Table 4 summarizes, GOcats extracts a notable number of uniquely enriched terms from the individual time point comparisons (UniqueEnrichedTerms_GOcats_). A few of these enriched terms (SupportedEnrichedTerms) are directly supported by the traditional method enrichment of the extreme time point comparisons.

**Table 4.**
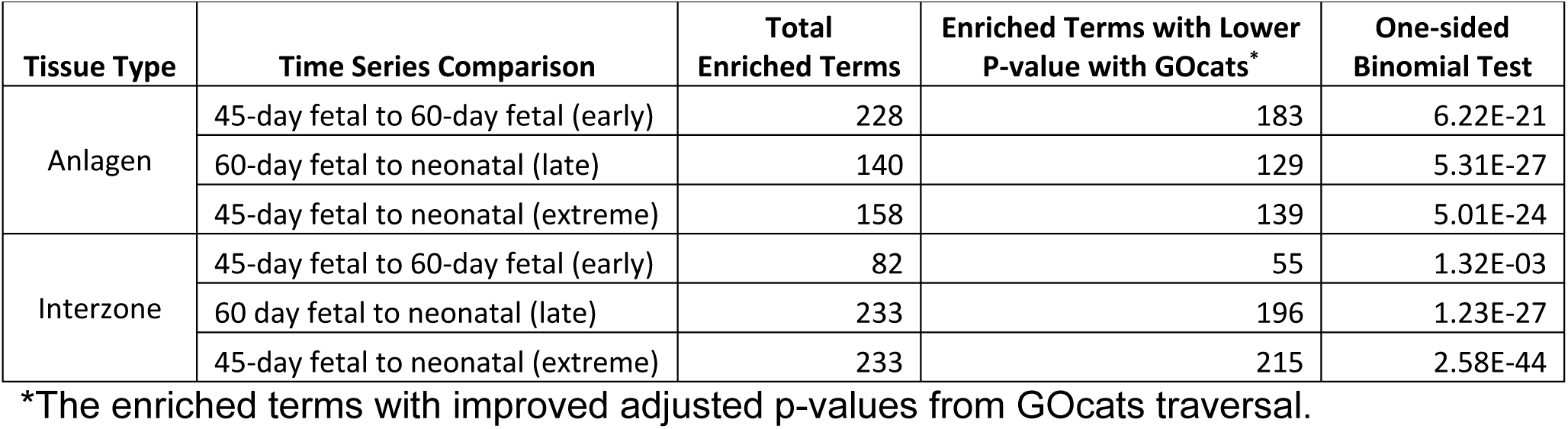
– Binomial test results for GOcats vs no_hp enrichment for horse cartilage development time point comparisons.

| Tissue Type | Time Series Comparison | Total Enriched Terms | Enriched Terms with Lower P-value with GOcats* | One-sided Binomial Test |
| --- | --- | --- | --- | --- |
| Anlagen | 45-day fetal to 60-day fetal (early) | 228 | 183 | 6.22E-21 |
|  | 60-day fetal to neonatal (late) | 140 | 129 | 5.31E-27 |
|  | 45-day fetal to neonatal (extreme) | 158 | 139 | 5.01E-24 |
| Interzone | 45-day fetal to 60-day fetal (early) | 82 | 55 | 1.32E-03 |
|  | 60 day fetal to neonatal (late) | 233 | 196 | 1.23E-27 |
|  | 45-day fetal to neonatal (extreme) | 233 | 215 | 2.58E-44 |

\*The enriched terms with improved adjusted p-values from GOcats traversal.

## Discussion

### Issues with semantic correspondence

As early as the late 1980s, explicit definitions of semantic correspondence for a relation between ontological terms have been stressed in the context of relational database design [19]. This includes concepts of part-whole (mereology), general-specific (hyponymy), feature-event, time-space (i.e spaciotemporal relations), and others. OBO’s and GO’s ontological edges are directional insofar as their relations accurately describe how the first node relates to the second node empirically, providing axioms for deriving direct semantic inferences. However, the directionality of these edges is ambiguous in that they do not explicitly describe how the terms relate to one another semantically in terms of scope, and this is due largely to the lack of explicit semantic correspondence qualifiers.

A simple way to avoid mapping problems associated with non-scoping relation direction is to omit those relations from analysis. This strategy avoids incorrect scoping interpretation at the expense of losing information. As an example, EMBL-EBI’s QuickGO term mapping service omits has_part type under its “filter annotations” by GO identifier options [16]. Furthermore, Bioconductor’s GO.db [20] also avoids mapping issues by indirectly omitting this relation; it uses a legacy MySQL dump version of GO which does not contain relation tables for has_part. We argue that while avoiding problematic relations altogether does avoid scope-specific mapping errors, it also limits the amount of information that can be gleaned from the ontology. By eliminating has_part from graphs created by GOcats, we see a ∼11% decrease in information content (as indicated by a decrease in the number possible mappings) in Cellular Component. Likewise, there is a 10% and 5% decrease of information content in Molecular Function and Biological Process, respectively (Table 2). Thus, omitting these relations from analyses removes a non-trivial amount of information that could be available for better interpretation of functional enrichment. However, the total impact is not completely appreciable here, because not all relations were evaluated in this study; only the scoping relations of is_a, part_of, and has_part. The potential for additional information loss is very high in Biological Process, for example, when considering the large number of unaccounted relations: regulates, positively_regulates, and negatively_regulates (Table 1). These relations add critical additional regulatory information to ontological graph paths, which would also be lost when ignoring the has_part relation, if they occurred along a path that also contained has_part. The same is also true for Molecular Function, although the prevalence of additional, non-scoping relations are lower.

Furthermore, automated summarization of annotations enriched in gene sets requires a more sophisticated evaluation of the scoping semantics contained in ontologies, which prior tools are not fully equipped to provide. M2S is one widely-utilized GO term categorization method that is available as part of the OWLTools Java application [21]. The Perl version of M2S has been integrated into the Blast2GO suite since 2008 [5, 22] and this gene function annotation tool has been cited in over 1500 peer-reviewed research articles (Google Scholar as of Nov. 28, 2017). We verified that the Perl and Java versions of M2S produced identical GO term mappings for a given dataset and GO slim, and therefore have the same mapping errors (Supplementary Data 2). Although the number of pM_F_s reported in the results represent the upper limit of the possible erroneous mappings, the fact that at least 120,000 of these exist in GO for the has_part relation alone or that the removal of this edge type results in up to an 11% reduction of information content provide bounds on the scope of the issue. To be clear, tools like M2S can be safe and not produce flawed mappings if they are used alongside ontologies that contain only those relations that are appropriate for evaluation, such as go-basic. However, we intentionally utilized *go-core* to illustrate the danger in using tools that do not provide explicit semantic control on how ontologies are utilized.

GOcats represents a step toward a more thorough evaluation of the semantics contained within ontologies by handling relations differently according to the linguistic correspondences that they represent. In the case of relations such as has_part, this involves augmenting the correspondence directionality when it is appropriate for the task at hand, which is to organize terms into categories. As a proof-of-concept, we classified the is_a, has_part, and part_of relations into a common “scoping” correspondence type and hard-coded assigned graph path tracing heuristics to ensure that they are all followed from the narrower-scope term to the broader-scope term. One caveat of this approach is that because of previously mentioned issues in universality logic, the inverse of has_part is not strictly part_of, but rather part_of_some. We argue that the unlikely misinterpretation of universality in this strategy is preferable to the loss of information experienced when using trimmed versions of ontologies for term categorization. To elaborate, most current situations calling for term categorization involve gene enrichment analyses. Spurious incorrect mappings through part_of_some edges would not enrich to statistical significance, unless a systematic error or bias is present in the annotations. Even if a hypothetical term categorization resulted in enrichment of a general concept that was not relevant to the system in question (i.e. “nucleus” enriched in a prokaryotic system), it would be relatively simple to reject such an assignment by manual curation and find the next most relevant term. Conversely, it is not reasonable to manually curate all possible missed term mappings resulting from the absence of an edge type in the ontology.

Another potential complication in semantic correspondence of relations is that some relations are *inherently* ambiguous. The clearest example of this again can be found in the well-utilized part_of relation. This relation is used to describe relations between physical entities and concepts (e.g. “nuclear envelope” part_of “endomembrane system”) and between two concepts (e.g. “exit from mitosis” part_of “mitotic nuclear division”) with no explicit distinction. To address the former issue, future work will augment our use of hard-coded categorization of semantic correspondences through the development of heuristic methods that identify and categorize these among the hundreds of relations in the Relations Ontology [2, 23]. As a good starting point, we suggest using five general categories of relational correspondence for reducing ambiguity (Table 1): scope (hyponym-hypernym), mereological, a subclass of scope (meronym-holonym), spatiotemporal (process-process, process-entity, entity-entity), active (actor-subject), and other.

### Using GOcats for annotation enrichment

While we reported the loss of information available for annotation enrichment with has_part excluded from GO and quantified the effect of incorrect inferences that can be made if has_part is included in GO during enrichment, these results only represent hypothetical effects that might be overcome when GOcats reinterprets this relation. One of GOcats’ original intended purposes was to improve the interpretation of results from annotation enrichment analyses. However, in the process of designing heuristics to appropriately categorize GO terminology, we also sought to overcome the limitations that come with following the traditional methods of path tracing along relations in GO. Here we focused on overcoming the loss of information encountered when ignoring has_part relations. Our solution was to re-evaluate these relations under the logic of part_of_some and invert the direction of has_part. While this re-interpretation is limited in usage, we believe that in the scope of annotation enrichment it is valid for reasons previously explained.

In our evaluation of enrichment results comparing GOcats ancestor paths to traditional GO ancestor paths in the enrichment analysis of a publicly-available breast cancer dataset, we demonstrate a highly statistically significant improvement (p=1.86E-25) in the statistical power of annotation enrichment analysis. Specifically, 182 out of 217 significantly enriched GO terms from the traditional analysis had improved p-values in the GOcats-enhance enrichment analysis. Moreover, we detect significantly enriched GO terms in the GOcats’ results that were not detected using the traditional analysis. The inclusion of the re-interpretation of has_part edges allowed for the significant enrichment (adjusted-p < 0.002 with FDR set to 0.01) of four terms related to purinergic nucleotide receptor signaling which has been implicated in predicting breast cancer metastasis in other studies [18]. We again confirmed this effect in our evaluation of GO annotation enrichment results of the horse cartilage development datasets. Here we saw an improvement in 67% to 92% of enriched terms across the six time point enrichment analyses. Fundamentally, the addition of part_of_some interpretation of has_part relations improves the statistical power of the annotation enrichment analysis, allowing the detection of additional enriched annotations with statistical significance from the same dataset. In addition, the GOcats annotation enrichment analysis extracts a notable number of uniquely enriched annotations from the neighboring, individual time point differential gene expression analyses. Some of these uniquely enriched terms are directly supported by the traditional annotation enrichment analysis of the extreme time point differential gene expression analyses (Table 4). These results on multiple datasets involving two separate experimental designs demonstrate the ability of utilizing GOcats-augmented ontology paths to derive additional information from annotation enrichment analyses.

To conclude, GOcats enables the simultaneous extraction and categorization of gene and gene product annotations from GO-utilizing knowledgebases in a manner that respects the semantic scope of relations between GO terms. It also allows the end-user to organize ontologies into user-defined biologically-meaningful concepts—a feature that we explore in-depth elsewhere [sister publication to cite]. This categorization lowers the bar for extracting useful information from exponentially growing scientific knowledgebases and repositories in a semantically safer manner. In summary, GOcats is a versatile software tool applicable to data mining, annotation enrichment analyses, ontology quality control, and knowledgebase-level evaluation and curation.

## Materials and Methods

Materials and methods are provided in Supplementary Data 1-5.

## Availability and Future Directions

The Python software package GOcats is an open-source project under the BSD-3 License and available from the GitHub repository https://github.com/MoseleyBioinformaticsLab/GOcats. Documentation can be found at http://gocats.readthedocs.io/en/latest/. All figures and supplementary data are available on the FigShare repository: https://figshare.com/s/952a4d001cc8850d6d5e along the code used to generate these results: https://figshare.com/s/9d55b2e5932992e6a068.

We are actively developing the codebase and appreciate any contributions and feedback provided by the community. We are extending the API and adding additional capabilities to handle more advanced annotation enrichment analysis use-cases.

## Acknowledgements

This work was supported in part by grants NSF 1252893 (Moseley), NIH 1U24DK097215-01A1 (Higashi, Fan, Lane, Moseley), and NIH UL1TR001998-01 (Kern).

